# Serum lipoproteins and lipoarabinomannan suppress the inflammatory response induced by the mycolactone toxin

**DOI:** 10.1101/2022.06.06.494992

**Authors:** Shailja Jakhar, Kiersten D. Lenz, Daniel E. Jacobsen, Philip A. Kocheril, Katja E. Klosterman, Harshini Mukundan, Jessica Z. Kubicek-Sutherland

## Abstract

*Mycobacterium ulcerans* is the causative agent of the chronic and debilitating neglected tropical disease Buruli ulcer (BU) which mostly affects children. The early detection and treatment of *M. ulcerans* infections can significantly minimize life-long disability resulting from surgical intervention. However, the disease is characterized by relatively few systemic systems as a result of complex host-pathogen interactions that have yet to be fully characterized, which has limited the development of both diagnostic and therapeutic approaches to treat BU. In this work, we study the interactions of the host immune system with two principle *M. ulcerans* virulence factors: mycolactone, an amphiphilic macrolide toxin, and lipoarabinomannan (LAM), a cell wall component of most mycobacterial pathogens. We observe that human lipoproteins have a profound effect on the interaction of both mycolactone and LAM with the immune system. Individually, both molecules are pro-inflammatory in the absence of serum and immunosuppressive in the presence of serum. When combined, mycolactone and LAM are immunosuppressive regardless of serum conditions. We also show that Toll-like receptor 2 (TLR2), a macrophage pathogen pattern recognition receptor, is critical for LAM immune stimulation but aids in mycolactone immunosuppression. These findings are a first step towards unraveling mycolactone-mediated immunosuppression during BU disease and may facilitate the development of effective diagnostics and therapeutics in the future.

**Author Summary:** Buruli ulcer (BU) is a neglected tropical disease caused by the pathogen *Mycobacterium ulcerans*. The principal virulence factors associated with it are the macrolide toxin mycolactone and the major cell wall component lipoarabinomannan (LAM). Here, we examine the impact of the amphiphilic biochemistry of mycolactone and LAM on their interaction with the human immune system. We show that both mycolactone and LAM associate with serum lipoproteins, and that this association is critical for the immune evasion seen in early-stage *M. ulcerans* infections. In the absence of serum, mycolactone is pro-inflammatory. Immunosuppression occurs only in the presence of human serum lipoproteins. In the presence of LAM, mycolactone is immunosuppressive, regardless of serum conditions. Immunosuppression is a hallmark of BU disease, and understanding the mechanisms of this immunosuppression can support the development of effective diagnostic and therapeutic strategies.

## 1. Introduction

Buruli ulcer (BU) is a neglected tropical disease caused by the pathogen *Mycobacterium ulcerans* and is characterized by painless skin lesions. The disease is particularly challenging to detect in early stages, since early symptoms are essentially non-existent and pain develops only later in infection (1-3). As a result, there has been a recent increase in BU cases in several areas, including Australia and Nigeria (4, 5). However, timeliness of therapeutic intervention can help to decrease the severity of the disease, since early stages of BU can be treated with antibiotics, but advanced stages require surgery and antibiotic therapy in a hospital (6-8).

The principal virulence factors associated with *M. ulcerans* are mycolactone, a secreted macrolide toxin, and lipoarabinomannan (LAM), a major cell wall component common to mycobacteria (2, 9, 10). Both virulence factors are amphiphilic, containing hydrophobic and hydrophilic moieties (11, 12), with a wide range of effects including cytotoxicity (13-15) and immunomodulation (10, 16, 17). Mycolactone’s immunosuppressive nature has led to the understanding that it is responsible for the painlessness of skin lesions in BU; however, recent findings have suggested that the course of disease is more complicated than originally proposed, and that the toxin demonstrates pro-inflammatory potential in later stages of infection (18). Similarly, the amphiphilic lipoglycan mannose-capped LAM, produced by pathogenic mycobacterial species, plays an important role in modulating key aspects of the host innate and adaptive immune responses (9). Both molecules are generally considered promising diagnostic biomarkers; mycolactone has been detected in blood prior to the onset of major symptoms of BU (19-21), and assays have been extensively developed for the detection of LAM in blood and urine, although these tests have low reported sensitivity (especially in patients living with comorbidities such as HIV) (22-25). The amphiphilic biochemistry of LAM and mycolactone makes them particularly challenging targets for diagnostics and therapeutics. In blood, these biomarkers associate with host lipoproteins, including high- and low-density lipoproteins (HDL and LDL) (12, 26-30), which limits the effectiveness of immunoassays targeting amphiphilic virulence factors. The interaction with human lipoproteins may also play an important role in mediating the immune response of the host during infection. The impact of both toxins’ amphiphilic biochemistry on their interaction with the host immune system has yet to be studied in depth.

The innate immune system has mechanisms for detecting pathogens using pattern recognition receptors (PRRs) against pathogen-associated molecular patterns (PAMPs) (31). PAMPs are conserved microbial virulence factors that are easily recognized by immune cell receptors, such as the Toll-like receptors (TLRs) (32). PAMPs are promising targets for both detection and anti-bacterial therapeutic strategies. Both LAM and mycolactone are recognized by multiple PRRs, including Toll-like receptor 2 (TLR2), which is primarily expressed on the surface of macrophages and dendritic cells during early infection (18, 33, 34). In the case of LAM, binding with immune receptors subsequently triggers the MyD88-dependent signaling pathway, leading to the production of pro- and anti-inflammatory cytokines (35, 36). Mycolactone has been shown to have opposing effects on the host immune system in early versus late stages of infection. In early infection, mycolactone targets the Sec61 channel to inhibit cytokine production and suppress the immune response (37-39). More recently, it has been shown that in later stages of infection, mycolactone induces the production of interleurkin-1β (IL-1β), a pro-inflammatory cytokine, in a TLR2-dependent mechanism (18). Mycolactone displays additional modes of action that cause cell damage and death, including binding to the Wiskott-Aldrich syndrome proteins to cause uncontrolled actin polymerization, and interaction with the pro-apoptotic regulator Bim (2, 40-42).

There have been inconsistent analyses on both LAM (35, 43-45) and mycolactone (18, 46-48) that suggest seemingly contradictory immune responses. The inconsistencies found could be due to the differential interaction of LAM and mycolactone with host lipoproteins, thereby making it difficult to interpret the biological significance of the findings. In this work, we investigate the binding of both LAM and mycolactone to serum lipoproteins, the implications these interactions have on the pathophysiology of BU, and the effect of this interaction on the nature of the host immune response.

## 2. Materials and Methods

### 2.1. Reagents and Materials

Dulbecco’s phosphate buffered saline (PBS, D8662) was obtained from Millipore Sigma (St. Louis, MO, USA). Synthetic mycolactone A/B stocks were a kind gift from the World Health Organization (Geneva, Switzerland) at 0.1 mg/mL in ethanol and stored at –20 °C protected from light until use (49). Purified lipoarabinomannan (LAM; *Mycobacterium tuberculosis*, strain H37Rv, NR-14848) was obtained through BEI Resources (NIAID, NIH). LAM was resuspended at 1 mg/mL in sterile nanopure water (Direct-Q 3 UV-R, Millipore Sigma). 60-μL aliquots were stored in low-retention tubes at –80 °C and thawed immediately prior to use. 1-mL aliquots of control human serum (BP2657100, Fisher Scientific, Hampton, NH, USA) were stored at −20 °C and thawed only once prior to use. 1-mL aliquots of de-lipidated human serum (J65516, BT-931, Alfa Aesar, Haverhill, MA, USA) were stored at −20 °C and thawed only once prior to use. Human high-density lipoproteins (HDL, L8039), low-density lipoproteins (LDL, L7914), very low-density lipoproteins (VLDL, LP1), and chylomicrons (CM, SRP6304) were purchased from Millipore Sigma and stored at 4 °C. Siliconized low-retention plastic microfuge tubes were used to store samples (02-681-320, Fisher Scientific).

### 2.2. Mycolactone-lipoprotein UV-vis Experiments

As described previously (12), mycolactone association with lipoproteins was quantified by UV-vis spectrophotometry (λ_max_ = 362 nm; log ε = 4.29). 1 μL of 100 μg/mL mycolactone A/B was added to 9 μL of lipoprotein solutions diluted in PBS (0, 0.125, 0.25, 0.5, or 0.75 mg/mL) and incubated at room temperature for 30 min to allow for mycolactone-lipoprotein association. UV-vis absorbance measurements were taken at 362 nm using 2 μL of each solution on a NanoDrop One^C^ (ND-ONEC-W, Thermo Fisher Scientific). Measurements were taken three times (technical replicates) and each experiment was repeated three times (biological replicates) for a total of nine absorbance measurements per lipoprotein concentration. Each lipoprotein concentration was measured in the absence of mycolactone and subtracted from the mycolactone measurement.

### 2.3. Tissue Lines and Reagents

The human monocyte cell line THP-1 (TIB-202) was purchased from the American Type Culture Collection (ATCC, Manassas, VA, USA). Human THP-1 TLR2 (-/-) cell line was custom-engineered by Horizon Inspired Cell Solutions (HD204-038, Horizon Discovery, Waterbeach, UK). RPMI-1640 medium (2 mM L-glutamine, 10 mM 4-(2-hydroxyethyl)-1-piperazineethanesulfonic acid, 1 mM sodium pyruvate, 4.5 g/L glucose, and 1.5 g/L sodium bicarbonate; 30-2001) and fetal bovine serum (FBS; 30-2020) were purchased from ATCC. 2-mercaptoethanol (M6250) and phorbol 12-myristate 13-acetate (PMA, P8139) were purchased from Millipore Sigma.

### 2.4. THP-1 Culture

THP-1 cells were revived in a T-25 Corning cell culture flask (CLS430639, Millipore Sigma) with RPMI-1640 supplemented with 0.05 mM 2-mercaptoethanol and 20% FBS. Once growth was established, cells were sub-cultured in T-75 tissue culture flasks with RPMI-1640 supplemented with 0.05 mM 2-mercaptoethanol and 20% FBS. THP-1 cells double every 24-48 h and were sub-cultured by addition of fresh growth media before reaching 10^6^ cells/mL.

### 2.5. THP-1 Adaptation to Low-serum Growth Conditions

Cells were adapted to growth in RPMI-1640 supplemented with 0.05 mM 2-mercaptoethanol containing 2.5% FBS in order assess the effect of adding serum and various lipoproteins on gene expression. Starting with 20% FBS upon cell revival, at every sub-culture the FBS concentration was reduced by half (20%, 10%, 5%, and 2.5%). RPMI-1640 containing 0.05 mM 2-mercaptoethanol and 2.5% FBS is referred to as low serum media.

### 2.6. Differentiation of THP-1 Monocytes to Macrophages

THP-1 cells were differentiated to macrophages using phorbol 12-myristate 13-acetate (PMA) by adapting previously published methods (50-52). Cells were first adapted to 2.5% FBS as described above, then sub-cultured in growth media containing 20 nM PMA. 1 mL of 8 × 10^4^ cells/mL was added to each well in a 24 well tissue culture plate (Corning Costar 07-200-740, Fisher Scientific) and incubated in 5 % CO_2_ at 37 °C for 24 h, at which point cells become adherent. After 24 h, PMA-containing media was removed, and cells were rinsed twice with growth media. 1 mL of growth media was added to each well and cells were allowed to differentiate for another 48–72 h in 5% CO_2_ at 37 °C. Differentiation was confirmed by flow cytometry as described previously (50).

### 2.7. LAM and Mycolactone THP-1 Exposure Experiments

LAM and mycolactone were pre-incubated with growth media containing 20% human serum, 2.5% human serum, 20% de-lipidated human serum, or human lipoproteins overnight at 4 °C to allow time for association. Lipoproteins were used at the following physiologically relevant concentrations: 0.5 mg/mL HDL (53), 1 mg/mL LDL (54), 0.15 mg/mL VLDL (55), 0.1 mg/mL CM (56). 0.5 mL of each solution was added to each well of differentiated THP-1 macrophages and incubated in 5% CO_2_ at 37 °C. After 24 h, the supernatant was removed, and cells were washed once with PBS (56064C, Millipore Sigma). RNA was extracted for gene expression analysis using the RNeasy Mini Kit (Qiagen, 74106) as per manufacturer instructions. RNA was pooled from at least four wells (technical replicates) to obtain enough RNA for gene expression analysis. Experiments were repeated at least three times (biological replicates).

### 2.8. Gene Expression Analysis by RT-PCR

cDNA synthesis was performed on the extracted RNA using the RT^2^ First Strand Kit (330404, Qiagen, Germantown, MD, USA). Initial gene expression profiling was performed using the 96-well Human Cytokines & Chemokines RT^2^ Profiler PCR Array (PAHS-150ZC, Qiagen) according to manufacturer instructions. Arrays contained 84 different human cytokines and chemokines, five housekeeping genes, and controls to monitor genomic DNA contamination, reverse transcription, and PCR efficiency. 96-well assays were used to down-select 18 genes of interest, which were ordered and tested on the remaining samples individually. PCR arrays and assays were run on an Applied Biosystems StepOnePlus thermocycler (Thermo Fisher Scientific) using RT^2^ SYBR Green ROX qPCR Mastermix (330529, Qiagen). Individual human RT^2^ primer assays and controls (330011): bone morphogenetic protein 2 (BMP2, PPH00549C), chemokine (C-C motif) ligand 1 (CCL1, PPH00701B), chemokine (C-C motif) ligand 2 (CCL2, PPH00192F), chemokine (C-C motif) ligand 20 (CCL20, PPH00564C), chemokine (C-C motif) ligand 3 (CCL3, PPH00566F), chemokine (C-C motif) ligand 5 (CCL5, PPH00703B), colony-stimulating factor 2 (CSF2, PPH00576C), colony-stimulating factor 3 (CSF3, PPH00723B), chemokine (C-X-C motif) ligand 1 (CXCL1, PPH00696C), chemokine (C-X-C motif) ligand 2 (CXCL2, PPH00552F), chemokine (C-X-C motif) ligand 5 (CXCL5, PPH00698B), chemokine (C-X-C motif) ligand 8 (CXCL8, PPH00568A), interleukin 1 alpha (IL-1α, PPH00690A), interleukin 1 beta (IL-1β, PPH00171C), interleukin 15 (IL-15, PPH00694B), interleukin 16 (IL-16, PPH00586A), interleukin 18 (IL-18, PPH00580C), tumor necrosis factor (TNF, PPH00341F), actin beta (ACTB, PPH00073G), hypoxanthine phosphoribosyltransferase 1 (HPRT1, PPH01018C), ribosomal protein large P0 (RPLP0, PPH21138F), reverse transcription control (RTC, PPX63340A), and genomic DNA contamination control (PA-031) were purchased from Qiagen. All PCR arrays and assays were subject to quality control monitoring for genomic DNA contamination, first strand synthesis (RTC), and real-time PCR efficiency (PPC, arrays only). Data were recorded as threshold cycle (C_T_).

### 2.9. Data analysis

UV-vis data analysis was performed with GraphPad Prism (v9.0.1, GraphPad Software, San Diego, CA, USA). One-way analysis of variance (ANOVA) testing was performed to assess differences between groups of data, with Tukey’s multiple comparisons testing performed post-hoc if a significant difference (*P* < 0.05) was identified. Gene expression analysis was performed using the Qiagen Gene Globe Data Portal (http://www.qiagen.com/geneglobe), by comparing test conditions (with LAM and/or mycolactone) to control conditions in the absence of these bacterial virulence factors. C_T_ values were normalized based on manual selection (arithmetic mean) using the housekeeping genes ACTB, HPRT1, RPLP0 and fold change/regulation was calculated with *P* values < 0.05 considered significant. Data were visualized with GraphPad Prism.

## 3. Results and Discussion

### 3.1. Mycolactone Directly Interacts with Serum Lipoproteins

Previous work from our laboratory and others has demonstrated a role for human lipoproteins in trafficking amphiphilic molecules in blood (27, 29, 57, 58). The interaction of LAM (26, 27) and mycolactone (12) with HDL and LDL has also been investigated. In this work, we studied the interaction of mycolactone with four major serum lipoproteins: HDL, LDL, VLDL, and CM. We tested each of these lipoproteins’ interaction with mycolactone at physiologically relevant concentrations (53, 54, 56, 59-61). We have previously described the interaction of LAM with serum lipoproteins (62).

Mycolactone is a small molecule with known low antigenicity (40). Because of a lack of available antibodies, reliable immunoassays have not been developed for the detection of mycolactone (40, 63). However, due to its conjugated π system, mycolactone has a UV-vis absorbance at 362 nm, allowing it to be easily measured in a spectrophotometer (12, 64, 65). To test mycolactone binding to serum lipoproteins, a range of lipoprotein concentrations between 0 and 0.75 mg/mL were incubated with 10 μg/mL mycolactone for 30 minutes, protected from light. Absorbance was then measured at 362 nm (**Figure 1a-d**). Background signal, measured as lipoprotein absorbance at 362 nm with no mycolactone, was subtracted from the absorbance measured with mycolactone-containing samples to correct for lipoprotein absorbance and scattering losses. Each serum lipoprotein showed the ability to bind to mycolactone with differing affinity. HDL and VLDL bound mycolactone with high affinity at lipoprotein concentrations as low as 0.125 mg/mL. At the physiologically relevant concentrations of 0.5 and 0.75 mg/mL (54, 61), insignificant but increased binding of mycolactone to LDL was also observed. The results for CM were inconclusive due to high background absorbance at 362 nm of the CM alone (**Table S1**). The results of our UV-vis experiments show that mycolactone interacts with native human serum lipoproteins with varying affinities. Understanding the interaction between these *M. ulcerans* virulence factors and serum lipoproteins is a critical step towards improving the diagnosis and treatment of BU.

**Figure 1.**
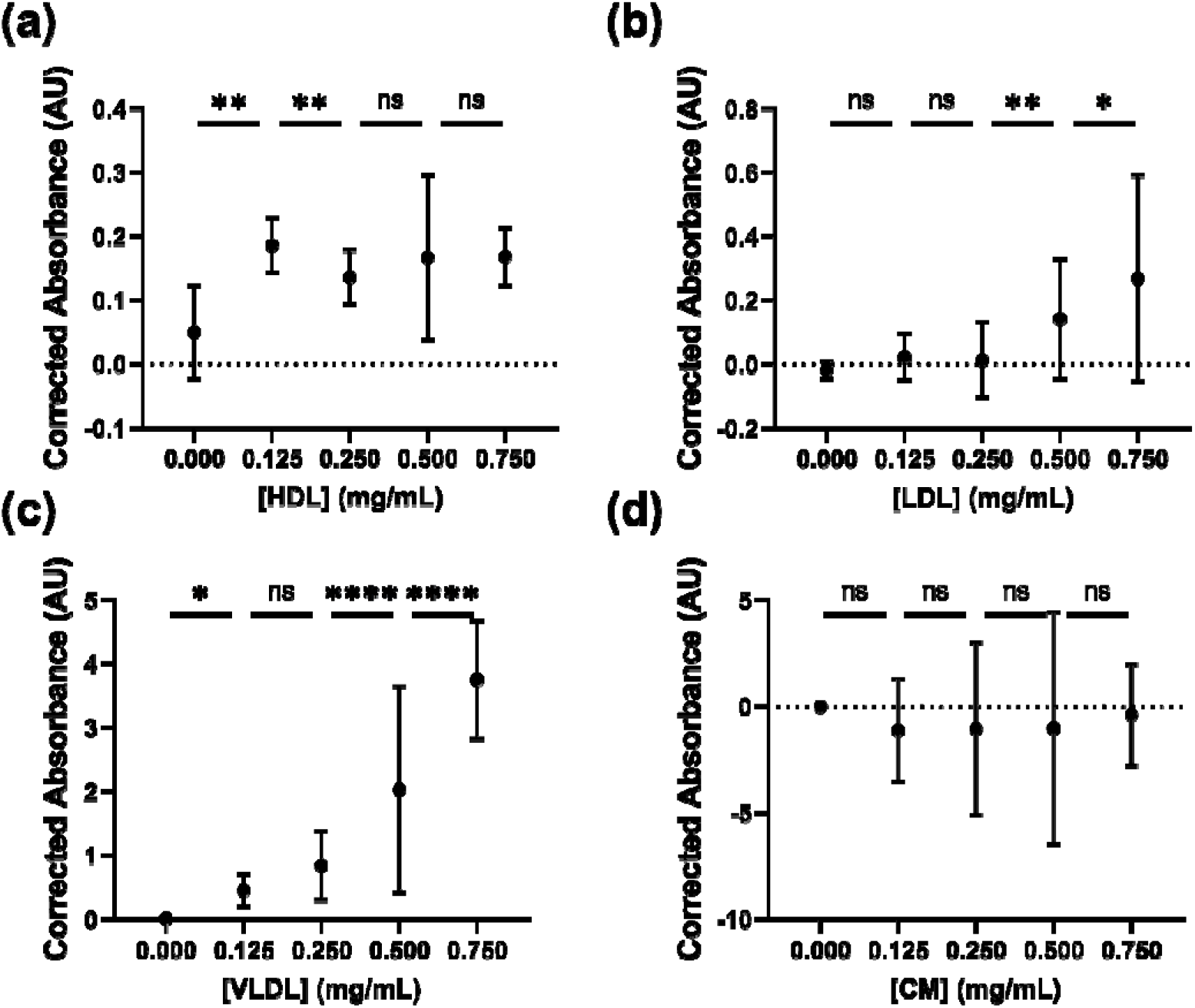
Mycolactone sequestration by serum lipoproteins. UV-vis absorbance measurements at 362 nm of 10 μg/mL mycolactone incubated with **(a)** HDL, **(b)** LDL, **(c)** VLDL, and (d) CM at concentrations from 0 to 0.75 mg/mL. Measurements (n = 9 replicates) are given as corrected absorbance at 362 nm (A_362_ mycolactone with lipoproteins – A_362_ lipoproteins alone at each respective concentration; AU, arbitrary units). Statistical significance was determined using a one-way ANOVA with a Tukey’s multiple comparisons test performed post-hoc if a significant difference was identified (**** P < 0.0001, ** P < 0.01, * P < 0.05; ns, not significant). Raw data and statistical analysis are provided in Table S1.

### 3.2. LAM is a Pro-inflammatory Virulence Factor in the Absence of Serum Lipoproteins

To examine the effect of individual serum lipoproteins on cytokine expression, THP-1 human macrophages were exposed to various conditions outlined in **Figure 2**. 10 μg/mL LAM was presented to THP-1 cells in culture media containing 1) 20% human serum, 2) 20% de-lipidated human serum (serum where lipoproteins are removed by centrifugation), 3) 2.5% human serum, and 4) individual human lipoproteins (HDL, LDL, VLDL, and CM). An exposure time of 24 hours and LAM concentration of 10 μg/mL were chosen based on the literature (44, 50-52, 66-71). For sustained growth in low serum conditions, we developed a protocol to adapt THP-1 cells to 2.5% serum as described in detail in the Methods section. Prior to LAM exposure, 20% serum and low serum adapted THP-1 cells were differentiated using PMA (50-52).

Initially, we examined the impact of LAM on the expression of 84 cytokines and chemokines using a high-throughput reverse transcription quantitative polymerase chain reaction (RT-qPCR) array (**Figure S1, Tables S2 and S3**). Prior to cell culture, 10 μg/mL of LAM was pre-incubated for 24 hours in just two media conditions (20% serum and low (2.5%) serum) in order to determine a sub-set of cytokines and chemokines that are affected by serum levels. The pre-incubation step allows for the association of LAM with serum lipoproteins prior to exposure to the cells in culture. Cell growth in the absence of LAM was used as a baseline for gene expression. It has been shown in the literature that LAM is an anti-inflammatory molecule that inhibits TNF and IL-12, while stimulating IL-10 production (72-75). However, our results indicate that this response is modulated by the lipoprotein concentration, suggesting a physiological range of outcomes. Indeed, we observed increased expression of TNF when cells were exposed to LAM in low serum as compared to LAM in the presence of 20% serum, indicating that the lipoproteins present in serum play a role in LAM’s ability to act as an anti-inflammatory agent. We did not observe changes in IL-12 and IL-10 between the different conditions. In response to LAM in the low serum condition, we observed high expression of CCL3, CXCL2, and CXCL8 that was not observed when lipoproteins in 20% serum were present. LAM in the absence of serum elicits a much greater cytokine and chemokine response compared to LAM in the presence of serum. This data indicates that serum lipoproteins suppress the interaction of LAM with the human immune system. These results were consistent between replicates (n = 3), and further studies were performed monitoring a subset of genes to validate and delve into this initial observation.

**Figure 2.**
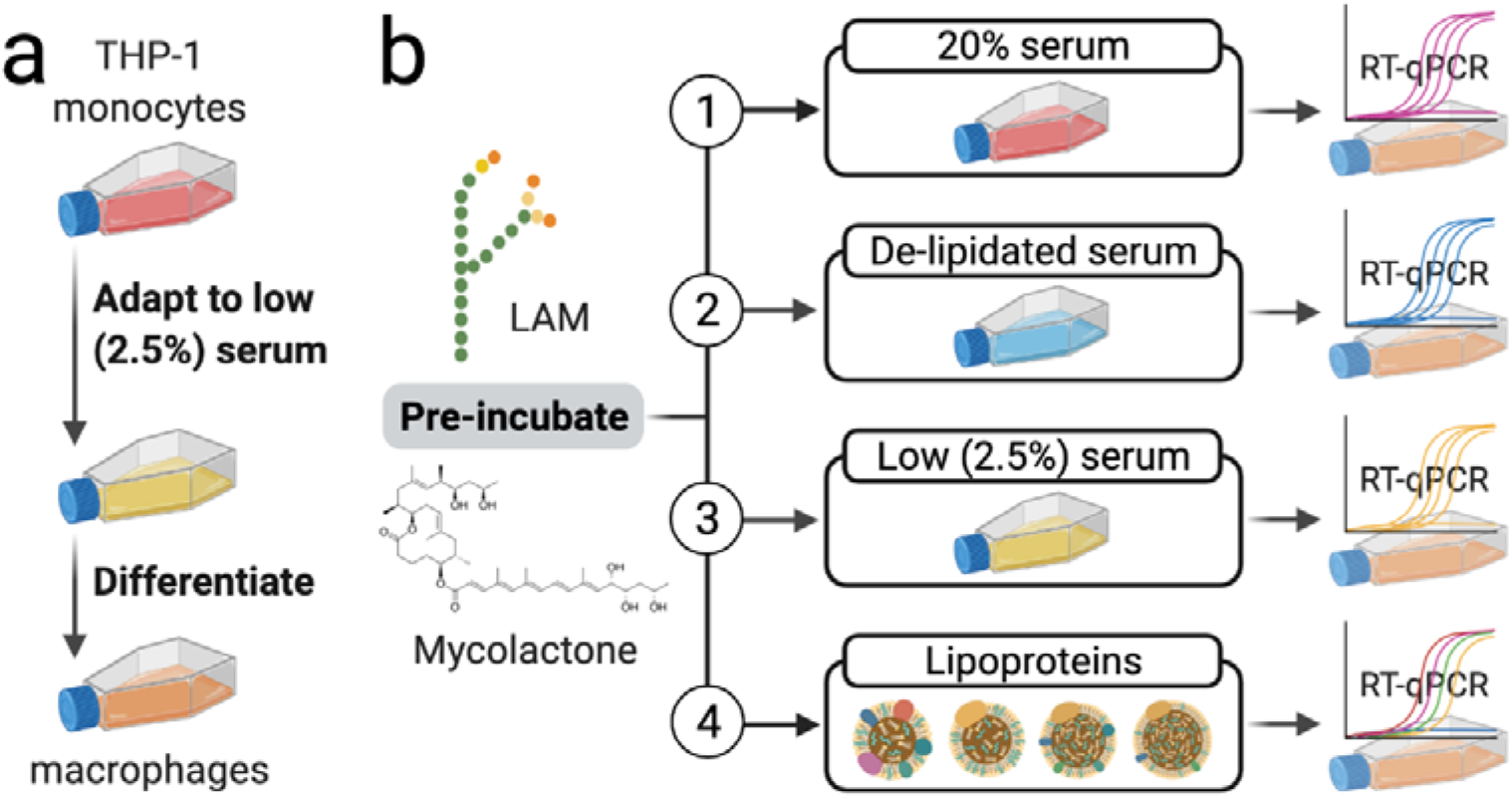
Overview of macrophage experiments. In these experiments, **(a)** THP-1 monocytes were adapted to growth in low serum conditions and then differentiated to macrophages. **(b)** Next, macrophages growing in low serum were exposed to LAM and mycolactone in the presence and absence of various serum and lipoprotein conditions. Created with BioRender.com.

### 3.3. Serum Lipoproteins Suppress the Pro-inflammatory Response Induced by LAM

To confirm the finding that the presence of lipoproteins in serum and their association with LAM impacts the cytokine and chemokine profile of human macrophage cells, we evaluated gene expression in 20% serum, 2.5% serum, de-lipidated serum, and 2.5% serum containing purified human lipoproteins (HDL, LDL, VLDL and CM), as outlined in **Figure 2**. LAM was allowed to associate with lipoproteins in the different conditions by pre-incubating overnight before exposure to THP-1 cells (**Figure 3**; see **Table S5** for significance testing results). LAM in de-lipidated serum and LAM in 2.5% serum conditions similarly showed pro-inflammatory responses with induction of CSF2, CCL3, CXCL2 and CXCL8. CCL3, also known as macrophage inflammatory protein (MIP-1α), is produced by a variety of cells including monocytes and macrophages and is involved in the acute inflammatory state (76). CXCL2, also known as MIP-2, and CXCL8, also known as IL-8, are associated with neutrophil infiltration in inflammatory diseases and attract macrophages and monocytes (77, 78). LAM in de-lipidated serum also shows an increase in IL-1β and TNF, which are chemotactic mediators involved in neutrophil recruitment to sites of inflammation (79). On the other hand, LAM in the presence of any individual lipoprotein shows suppression of this immune response. Interestingly, VLDL and CM showed higher suppression of cytokine signaling than HDL and LDL. This trend could be due to the larger particle size of VLDL (30-90 nm) and CM (200-600 nm) as compared to HDL (7-13 nm) and LDL (21-27 nm), which might provide them with greater surface area to associate with LAM (80-82). Additionally, the different molar concentrations of the various serum lipoproteins used in these experiments (based on normal physiological concentrations) could further contribute to the observed differences in cytokine signaling upon exposure to LAM.

**Figure 3.**
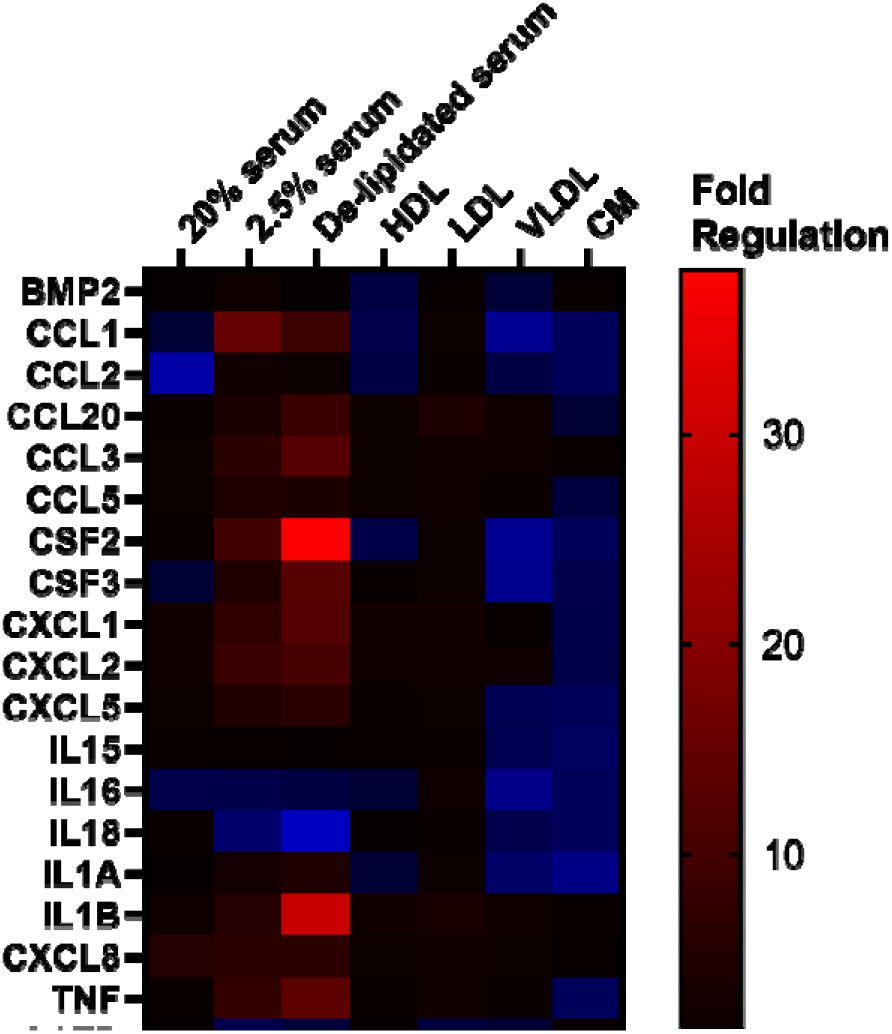
LAM is pro-inflammatory in the absence of serum lipoproteins. Cytokine profiles of THP-1-derived macrophages exposed to LAM at 10 μg/mL in media containing 20% serum, 2.5% serum, de-lipidated serum, and 2.5% serum containing the individual lipoproteins HDL, LDL, VLDL, and CM. The housekeeping genes ACTB, HPRT1, RPLP0 were used for data normalization. Values are plotted as fold regulation from 3 independent experiments (n = 3). See Tables S4 and S5 for raw C_T_ and Fold Regulation data, respectively.

### 3.4. Mycolactone is Pro-inflammatory in the Absence of LAM and Serum Lipoproteins

After observing that mycolactone and LAM interact directly with serum lipoproteins, we investigated the effects of both molecules on the cellular immune response by exposing THP-1 cells to 3 ng/mL mycolactone and 10 μg/mL LAM in 20% serum and low (2.5%) serum conditions (**Figure 4**; see **Table S5** for significance testing results).

**Figure 4.**
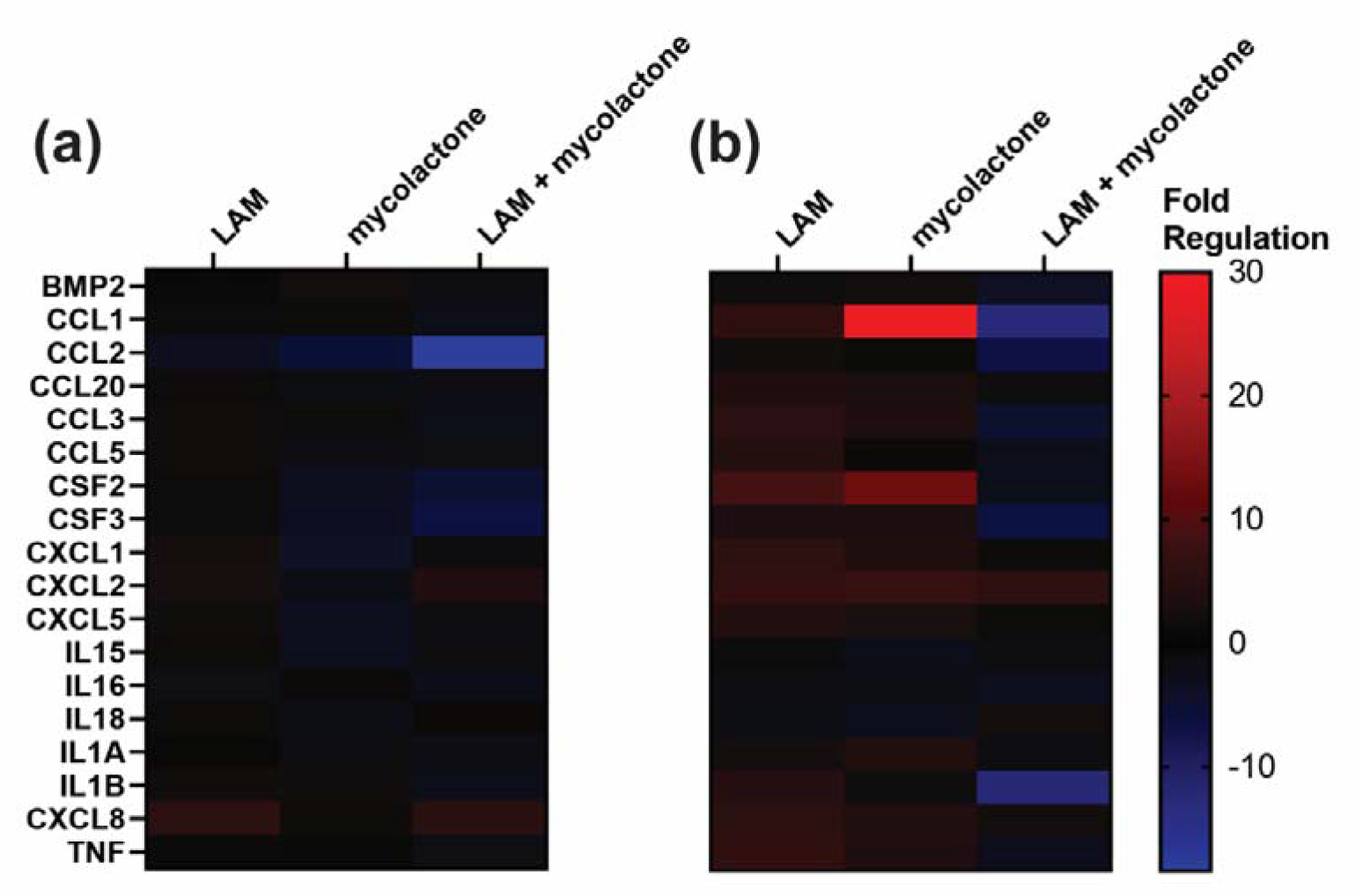
Immunosuppression occurs when mycolactone and LAM are co-administered. Cytokine profiles of THP-1-derived macrophages exposed to 10 μg/mL LAM, 3 ng/mL mycolactone, and both mycolactone and LAM co-administered in **(a)** 20% serum and **(b)** 2.5% serum conditions. See Tables S4 and S5 for raw C_T_ and Fold Regulation data, respectively.

In media containing 20% serum, effectively no pro-inflammatory activation was observed in cells exposed to LAM, mycolactone, or both (**Figure 4a**). In contrast, in media with 2.5% serum, both LAM and mycolactone are pro-inflammatory, suggesting that the lipoproteins present in serum play an important role in suppressing the immune response of macrophages exposed to LAM or mycolactone. Interestingly, cells exposed to LAM and mycolactone together displayed little to no cytokine response whether serum was present or not, suggesting that LAM and mycolactone act together to suppress the immune system.

### 3.5. TLR2 Inhibits Mycolactone Stimulation of the Immune System

To determine the role of TLR2 in the cellular response to LAM and mycolactone, cytokine expression of TLR2 wild-type (+/+) cells was compared to TLR2 knockout (-/-) cells (**Figure 5**; see **Table S5** for significance testing results). TLR2 has been reported as a down-stream effector of both mycolactone (18, 83) and LAM (33, 44, 71, 84). The presence of 20% serum suppressed the cytokine response to LAM regardless of TLR2 status (**Figure 5a**). In the absence of serum and presence of TLR2, LAM is pro-inflammatory. This response is sup-pressed in the absence of TLR2 (-/-), which suggests that LAM signaling is mediated by TLR2 as described previously (33, 44, 71, 84). In the absence of both TLR2 and serum, LAM exhibits effectively neither prornor anti-inflammatory activation, suggesting that TLR2 and serum constituents exclusively mediate LAM-induced immune responses.

**Figure 5.**
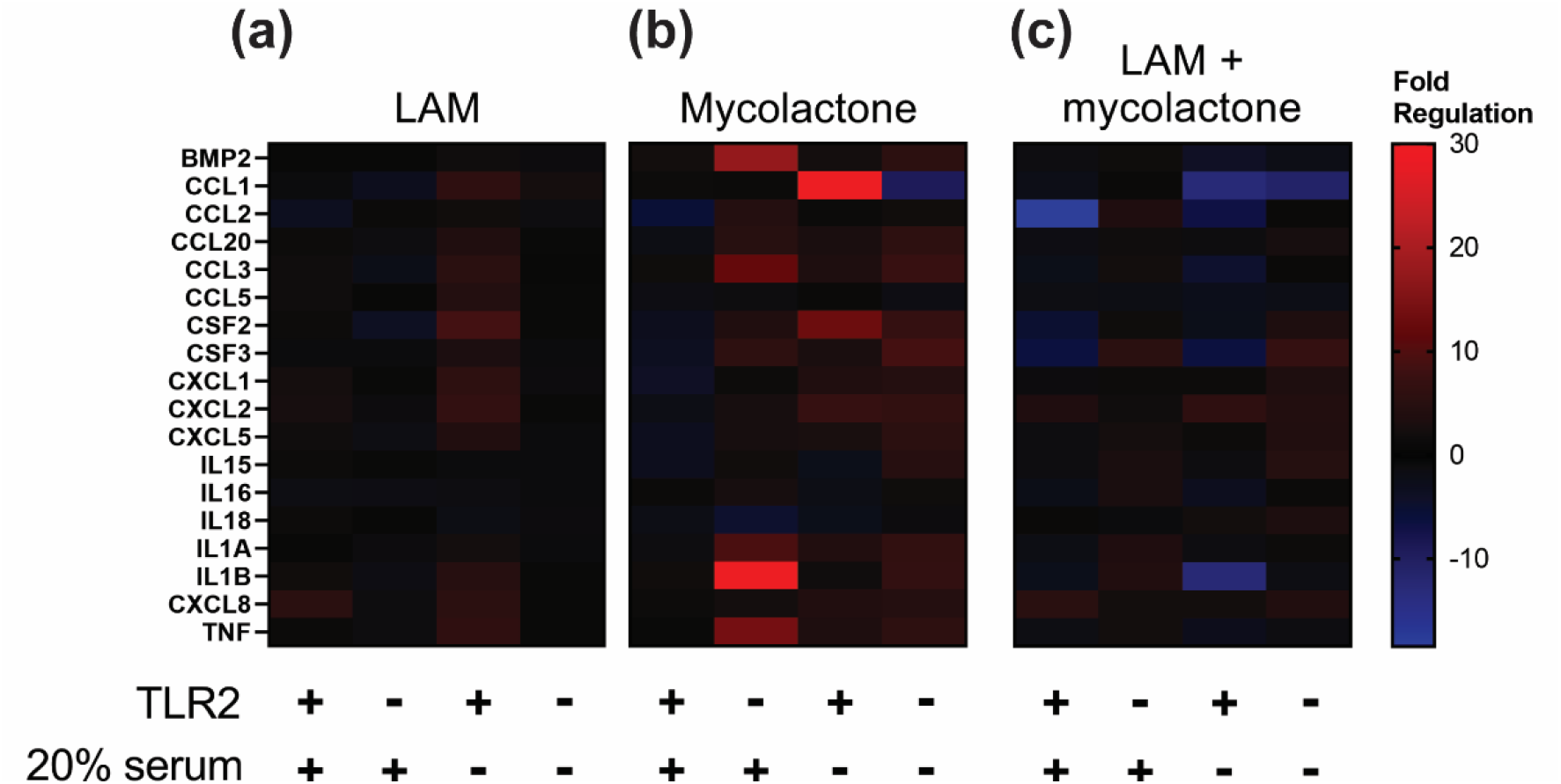
TLR2 inhibits mycolactone stimulation of the immune system. Cytokine profiles of THP-1-derived macrophages containing wild-type TLR2 (+/+) or TLR2 knockout (-/-) exposed to **(a)** 10 μg/mL LAM, **(b)** 3 ng/mL mycolactone, and **(c)** both LAM and mycolactone co-administered. See Tables S4 and S5 for raw C_T_ and Fold Regulation data, respectively.

TLR2 plays a different role in the immune response to mycolactone (**Figure 5b**). In the absence of TLR2 (-/-), mycolactone was pro-inflammatory regardless of whether serum was present or not. Similarly, in the absence of serum, mycolactone was pro-inflammatory rergardless of whether TLR2 was present or not. Mycolactone only failed to elicit a pro-inflammatory response when both TLR2 and serum were present. These results indicate that mycolactone is pro-inflammatory, but TLR2 and serum together inhibit its stimulation of the immune system.

When mycolactone and LAM were co-administered, we observed limited immune rersponses (**Figure 5c**). However, gene expression in the TLR2 knockout (-/-) cell line was sigrnificantly higher than the gene expression in wild-type TLR2 (+/+) in both 2.5% serum (P < 0.001) and 20% serum (P < 0.01) conditions. These data corroborate our previous finding that TLR2 inhibits mycolactone’s stimulation of the immune system. In the absence of TLR2 (-/-), co-administration of LAM and mycolactone resulted in immune induction, although the effect was reduced when compared to mycolactone alone. This finding indicates that LAM is immunosuppressive in the absence of TLR2. Finally, the presence of serum suppresses the immune response with and without TLR2.

### 3.6. LAM and Mycolactone May Lead to Pro-inflammatory Activation through HDL Dysfunction and Membrane Disruption

The association of LAM and mycolactone with serum lipoproteins provides insights into the pathophysiology of BU. For example, as we noted in the Introduction, mycolactone is thought to be immunosuppressive in the early stages of BU and pro-inflammatory in the late stages of BU (18). Our results indicate that the immunosuppressive nature of mycolactone is mediated by serum lipoproteins and TLR2, likely involving direct association with serum lipoproteins. Notably, serum lipoproteins do not have unlimited capacity for toxins that have been sequestered from the blood, and saturation of lipoproteins with toxins has the potential to change their biological activity. Similar effects have been observed in chronic kidney disease; saturation of HDL with symmetric dimethylarginine, a circulating kidney biomarker, renders the HDL dysfunctional (85). The dysfunctional HDL then initiates TLR2 signaling on endothelial cells, leading to a variety of deleterious effects, including hypertension and increased superoxide production (86).

Therefore, in the early stages of BU, mycolactone could be entirely sequestered by HDL, leading to immunosuppression. But in the later stages of BU, circulating HDL could become saturated with mycolactone, leading to pro-inflammatory activation. It is worth noting that saturated, dysfunctional lipoproteins may not be involved in pro-inflammatory activation in BU, but instead newly produced mycolactone that is unable to be sequestered may begin to circulate at appreciable concentrations in the bloodstream, leading to pro-inflammatory activation by some other mechanism. Similarly, LAM’s TLR2-dependent immune stimulation, which we observed in the absence of serum lipoproteins, may also exhibit lipoprotein saturation behavior. Further biophysical characterization of the interactions between serum lipoproteins and both LAM and mycolactone will be invaluable in deciphering the interaction of *M. ulcerans* with the immune system.

The identities of the upregulated pro-inflammatory cytokines that we measured by RT-qPCR can provide further insight into the specific biochemical pathways activated in BU. Using the Kyoto Encyclopedia for Genes and Genomes (KEGG), we searched for pathways involving the most significantly upregulated cytokines (87-89). We noticed that a handful of our most significantly upregulated cytokines (TNF, IL-1β, CSF2, and IL-8) were noted as downstream of membrane disruption in shigellosis through the NF-κβ signaling pathway (KEGG: map05131) (87-89). Given the amphiphilic natures of LAM and mycolactone, membrane disruption provides a plausible mechanistic basis for immune stimulation in late-stage BU.

## 4. Conclusions

In this manuscript we have identified an important relationship between the immuno-suppressive effects of mycolactone and LAM and the presence of serum lipoproteins. The absence of serum results in a cytokine response from macrophages exposed to either virulence factor. This response indicates that amphiphilic PAMPs avoid immune stimulation by associrating themselves with lipoproteins found in serum. We additionally showed that both LAM (62) and mycolactone can interact with any of the four major serum lipoprotein classes, indircating that this association is nonspecific and likely due to a combination of hydrophobic and hydrophilic interactions. Interestingly, the presence of both molecules is immunosuppressive with or without serum, indicating that the two molecules inhibit macrophages along different, but not unrelated, pathways. Understanding the interplay between these molecules and serum lipoproteins will be integral to improving the diagnosis and treatment of *Mycobacterrium*-associated illness, including BU. Based on these findings, another implication to consider is that human lipoprotein concentrations are impacted by a variety of metabolic and physiological processes such as age, diet, co-morbidities (obesity, dyslipidemia, cardiovascular disease, diabetes), and co-infections (HIV, malaria) (90-94). Thus, the susceptibility of a given individual to BU is a complex interplay of the pathogen with host health and physiology.

We have also investigated the interaction between mycolactone, LAM, and TLR2. When TLR2 was knocked out, mycolactone induced an immune response in macrophages regardless of the presence or absence of serum. This response indicates that TLR2 plays an inhibitory role in mycolactone immune stimulation. In the presence of both LAM and mycolactone, as during an *M. ulcerans* infection, TLR2 knockout induces a small but significant cytokine response. LAM dampens the mycolactone-dependent immune stimulation that occurs in the absence of TLR2, suggesting that LAM and mycolactone each play immunosuppressive roles during infection.

Many bacterial PAMPs are amphiphilic and are transported within the body by host lipoproteins. These lipoprotein interactions help the amphiphilic PAMPs escape detection by the immune cells of the host. Here, we have examined the potential of both LAM and mycolactone to directly interact with human lipoproteins, as well as the effect this interaction has on immune signaling. We have also characterized mycolactone and LAM’s interactions with TLR2, including that mycolactone acts with TLR2 in an immunosuppressive fashion. As they are prominent molecules in the pathogenesis of BU, a better understanding of the activities of both LAM and mycolactone will remain important in designing more effective diagnostic and therapeutic strategies.

## Supplementary Materials

The following supporting information is available; Table S1: Raw UV-vis data and statistical analysis of mycolactone interaction with human lipoproteins; Table S2: Raw RT-PCR CT values from 96-well human cytokine and chemokine arrays for THP-1 experiments with LAM. Table S3: Fold regulation from 96-well human cytokine and chemokine arrays for THP-1 experiments with LAM. Table S4: Raw RT-PCR C_T_ values from THP-1 and TLR(-/-) experiments with LAM and mycolactone; Table S5: Fold regulation and *P*-values from THP-1 and TLR2(-/-) experiments with LAM and mycolactone.

## Author Contributions

Conceptualization, J.K.S.; methodology, S.J. and J.K.S.; validation, S.J.; formal analysis, K.D.L., D.E.J., and J.K.S.; investigation, S.J., K.D.L, K.E.K., and J.K.S.; resources, H.M. and J.K.S.; writing—original draft preparation, S.J., K.D.L., D.E.J., P.A.K., and J.K.S.; writing—review and editing, S.J., K.D.L., D.E.J., P.A.K., K.E.K., H.M., and J.K.S.; visualization, K.D.L., D.E.J., and J.K.S.; supervision, H.M. and J.K.S.; project administration, H.M. and J.K.S.; funding acquisition, H.M. and J.K.S. All authors have read and agreed to the published version of the manuscript.

## Institutional Review Board Statement

Not applicable.

## Informed Consent Statement

Not applicable.

## Data Availability Statement

Ra data are available in the Supplemental Information (Tables S1-S5).

## Acknowledgments

We are grateful to the World Health Organization for their kind donation of synthetic mycolactone used in this study. We also thank Dr. Eric Nuermberger, Dr. Paul Converse, Dr. Basil I. Swanson, Dr. Susan E. Dorman, Dr. Dominique N. Price, and Dr. Loreen R. Stromberg for technical guidance and helpful discussions during the course of this work. This work was performed at the Los Alamos National Laboratory, which is operated by Triad National Security, LLC, for the National Nuclear Security Administration of U.S. Department of Energy (Contract No. 89233218CNA000001). The views expressed in this article are those of the authors and do not reflect the official policy or position of the U.S. Government.

