## Supplementary material for "Serum lipoproteins and lipoarabinomannan suppress the inflammatory response induced by the mycolactone toxin": Figure S1

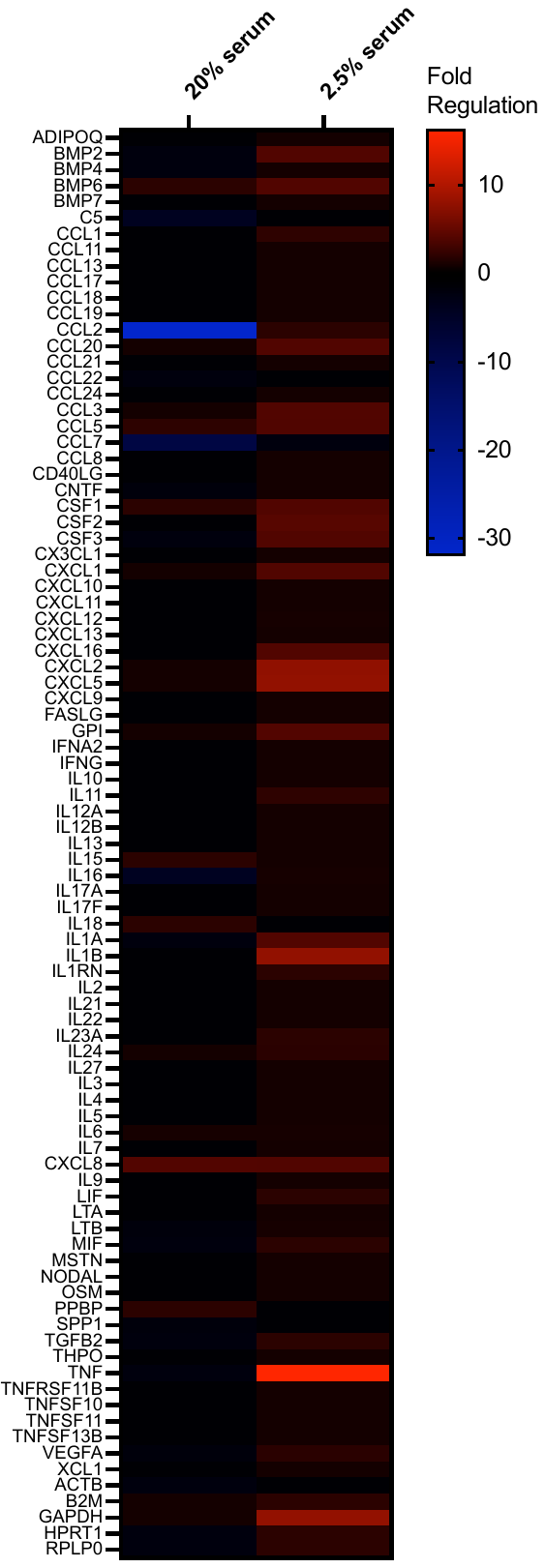
**Figure S1.** **LAM is a pro-inflammatory virulence factor in the absence of serum lipoproteins.** The heat map shows fold regulation in gene expression when 10 µg/mL LAM is incubated overnight with 20% serum versus low (2.5%) serum media comparing cells exposed to LAM to those incubated in the same conditions without LAM. These 96-well arrays were used to down-select genes for targeting in further experiments. See Tables S2 and S3 for raw C_T_ and Fold Regulation data, respectively.
